# NetworkCommons: bridging data, knowledge and methods to build and evaluate context-specific biological networks

**DOI:** 10.1101/2024.11.22.624823

**Authors:** Victor Paton, Denes Türei, Olga Ivanova, Sophia Müller-Dott, Pablo Rodriguez-Mier, Veronica Venafra, Livia Perfetto, Martin Garrido-Rodriguez, Julio Saez-Rodriguez

## Abstract

**Summary:** We present NetworkCommons, a platform for integrating prior knowledge, omics data, and network inference methods, facilitating their usage and evaluation. NetworkCommons aims to be an infrastructure for the network biology community that supports the development of better methods and benchmarks, by enhancing interoperability and integration.

**Availability and Implementation:** NetworkCommons is implemented in Python and offers programmatic access to multiple omics datasets, network inference methods, and benchmarking setups. It is a free software, available at https://github.com/saezlab/networkcommons.

**Supplementary Data:** Contribution guidelines, additional figures, and descriptions for data, knowledge, methods, evaluation strategies and their implementation are available in the Supplementary Data and in the NetworkCommons documentation at https://networkcommons.readthedocs.io/.

## 1. Introduction

Network biology leverages computational methods to clarify complex interactions among molecules, including genes, proteins, and metabolites (Zitnik et al. 2023). In particular, context-specific network inference methods integrate context-agnostic prior knowledge with omics data to identify subnetworks associated with specific conditions, such as perturbations or diseases (Garrido-Rodriguez et al. 2022). These networks then enable various downstream applications, including omics data interpretation and identification of potential drug targets (Perrone et al. 2024).

A wide range of tools have been developed to infer networks from different combinations of knowledge and data. However, these tools do not share a common application programming interface (API). This lack of interoperability hinders benchmarking of these methods as well as the reuse and combination of their modules.

Here, we present NetworkCommons, a Python package that unifies programmatic access to (1) prior knowledge, (2) omics data, and (3) network contextualization methods. With these components, we provide a platform for users to create, apply and evaluate custom subnetwork identification problems. The value of this platform is illustrated by providing access to four preprocessed omics datasets, eight network inference methods, and four benchmarking setups, as a stepping stone towards a ‘Network commons’, an ecosystem of resources and methods for network biology. All code and extensive documentation are available at https://networkcommons.readthedocs.io/.

## 2. Results

NetworkCommons offers a high-level API for accessing prior knowledge, omics data, and contextualization methods, allowing users to integrate, evaluate, and visualize generated subnetworks. The package is divided into four primary modules (Figure 1).

**Figure 1:**
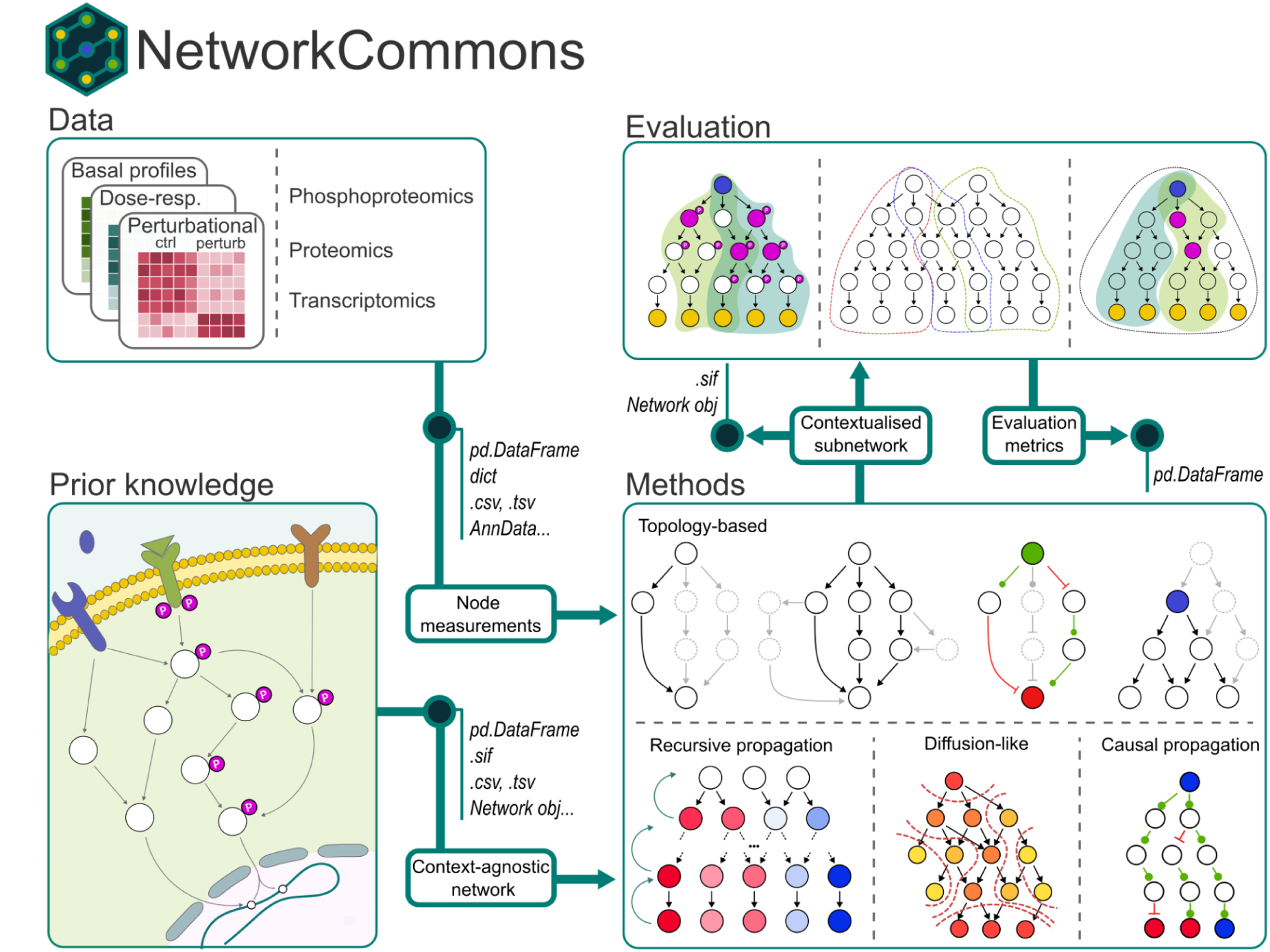
Overview of NetworkCommons. The package connects four different areas: Data, Prior Knowledge), Contextualization Methods, and Evaluation.

The first module provides access to *prior knowledge networks* (PKNs), representing various types of context-independent, protein-level interactions. Users can either use one of the built-in PKNs or import their own by converting it into a NetworkCommons PKN object (Supp. Data IV.1.). The currently built-in PKNs include only directed networks. Directed networks are of particular interest as they contain the minimal molecular information needed to generate causal hypotheses and statements (Touré et al. 2021). Of note, the API and some of the included methods are also fully compatible with undirected graphs. The networks may also include a *weight* interaction attribute to indicate up- or down-regulation. These weights can also convey other quantitative or qualitative information, although most of the network contextualization methods do not directly employ this information. The built-in prior knowledge relies primarily on *OmniPath* (Türei et al. 2021), a database combining dozens of resources, covering protein-protein, kinase-substrate, ligand-receptor interactions, as well as gene regulatory networks. Other sources can be easily added (Supp. Data IV.1.).

The *omics data* module provides built-in access to preprocessed omics datasets, encompassing diverse types of omics data across multiple biological contexts. We illustrate the access to four datasets, and also how users can connect their own datasets to the platform (Supp. Data IV.2.). All datasets are managed through *pandas* data frames, with an additional interface available for *AnnData* objects from the *scverse* framework (Virshup et al. 2023), as illustrated in a dedicated vignette that employs perturbational data from *pertpy (Heumos et al. 2024)*. The first among the datasets is dose-response phosphoproteomics data from *DecryptM,* as preprocessed by its authors (Zecha et al. 2023). Second, we curated and reprocessed data from the *PANACEA DREAM challenge* (Douglass et al. 2022), which profiles single-dose transcriptomic responses to 32 kinase inhibitors across 11 cancer cell lines. Third, we added basal metabolomics and transcriptomics data from the *NCI-60* cancer cell line collection (Shoemaker 2006). Lastly, we provide access to harmonized multi-omics data from the *Clinical Proteomic Tumor Analysis Consortium* (Li et al. 2023), which includes genomics, transcriptomics, proteomics, and phosphoproteomics data for various tumor and adjacent normal tissue samples across a wide range of cancer types. This second module offers access to a variety of large datasets spanning basal, perturbed, and clinical contexts, and users are welcome to input their own datasets.

The *methods* module offers a consistent API for various network contextualization methods. Each method requires, at a minimum, a PKN object and a set of upstream and downstream measurements of molecular activities. The output of each method is a contextualized subnetwork. NetworkCommons already includes a range of contextualization methods, such as topology-based, recursive propagation, diffusion approaches, and causal propagation (described in detail in Supp. Data II.). New methods can be included from external modules by wrappers, or implemented directly within NetworkCommons, using libraries already supported in the framework, such as *NetworkX* (Hagberg, Schult, and Swart 2008) or *CORNETO* (Rodriguez-Mier et al. 2024). For example, the causal reasoning method *CARNIVAL* (Liu et al. 2019), as implemented in *CORNETO,* is made available in NetworkCommons by a minimal wrapper (Supp. Data II.6.). We also incorporate SignalingProfiler (Venafra, Sacco, and Perfetto 2024), which integrates topology-based methods and *CORNETO* to generate hierarchical signaling models (Supp. Data II.7.). NetworkCommons also provides network visualization and other plotting functions to support quality control, data exploration and result interpretation.

The fourth main component of NetworkCommons, the *evaluation* module, allows for the benchmarking of contextualization methods. We illustrate this with four strategies to evaluate context specific subnetworks, using the datasets already available via the *omics data* module. Each evaluation strategy requires context specific networks as input, and outputs a set of evaluation metrics. The first evaluation strategy makes use of the perturbational profiles from compounds in the *PANACEA* project. Here, the networks are scored according to the number of recovered off-target proteins. The second strategy uses *DecryptM* data to recover phosphorylation changes based on sensitivity to drug perturbations. The third strategy scores subnetworks based on the enrichment of an *a priori* expected gene set. Lastly, we used harmonized data from the *Clinical Proteomic Tumor Analysis Consortium* to showcase our fourth evaluation strategy. Here, we employed protein abundance from proteomics, and activity scores inferred from transcriptomics to contextualize networks. These are then evaluated based on the enrichment of kinase activities from phosphoproteomics. More details about the strategies are available in Supp. Data III., about the implementation and creating new strategies, see Supp. Data IV.4.

## 3. Discussion

In this note, we present NetworkCommons, a platform for the integration of omics data and prior knowledge with methods to generate context-specific molecular interaction networks. With our design, we aim for interoperability, ease of access and robustness of existing and future components. While the current version represents the beginning of a broader effort, we already illustrate its potential by providing integrated access to several data sources, methods, and different evaluation strategies for contextualization methods. We propose a programmatic environment where these evaluation strategies can be tested. We provide four of them to demonstrate this concept, and as a starting point for method developers to evaluate new approaches.

In the future, we aim to expand the ecosystem to include more datasets, methods and prior knowledge, to provide support for a wider range of applications. We plan to incorporate additional methods, such as *phuEGO* (Giudice et al. 2024), and expand our methods towards, for example, undirected networks and genetic data (Barrio-Hernandez et al. 2023), or functional and disease annotations, leveraging integrated knowledge graphs (Lobentanzer et al. 2023). We also aim to further develop the API by introducing higher level objects, expanding the use of *AnnData* for handling omics data (Virshup et al. 2023), open ways towards Boolean modeling (Ruscone et al. 2024), and improve the connection with other ecosystems such as *scverse* for single-cell data analysis and preprocessing, and in particular *pertpy* to leverage perturbational data (Heumos et al. 2024).

With NetworkCommons, we aim to create a meeting point for the network biology community. We envision it as a key infrastructure, modules and utilities that everyone can reuse for their own developments and applications, and supports a broader collective effort to build robust and efficient context-specific network inference methods and advanced benchmarks. To facilitate this, we created contribution guidelines, inviting current and future developers within the network biology field to contribute with data, prior knowledge, methods and evaluation strategies. By providing an unified platform for data, prior knowledge and methods, we aim not only to empower users to use mechanistic network approaches in their research, but also to encourage the community to join efforts and push the boundaries of network biology.

## 4. Supplementary data

### I. Omics data and prior knowledge

#### I.1. Perturbational transcriptomics data from Douglass et al

This dataset contains 1728 RNA-Seq profiles of 11 cell-lines perturbed with 32 kinase inhibitors. Cell-lines were treated at a 48-hour IC20 for each drug and RNA was collected for sequencing at 24 hours, 2 replicates for each drug-cell-line pair (Douglass et al. 2022). Originally, this resource served as the basis for a *DREAM Challenge* assessing the accuracy and sensitivity of computational algorithms for de novo drug polypharmacology predictions. NetworkCommons provides raw files for count data and metadata, as retrieved in the original page. In addition, differential expression and TF activity tables are provided.

The differential expression statistics were obtained via *FLOP* (Paton et al. 2024), using *FilterbyExpr* and *DESeq2,* one of the top performer combinations in the benchmarking study. The contrasts were set, per cell line, between each drug and the DMSO control. The TF activity tables were obtained also via *FLOP,* using univariate linear models as implemented in decoupler. Only contrasts with more than 30 differentially expressed genes underwent downstream TF enrichment.

#### I.2. Dose-response perturbational phosphoproteomics from Zecha et al

This dataset contains the profiling of 31 cancer drugs in 13 human cancer cell line models, resulting in 1.8 million dose-response curves. The data includes 47,502 regulated phosphopeptides, 7,316 ubiquitinated peptides, and 546 regulated acetylated peptides (Zecha et al. 2023). NetworkCommons provides the files containing EC50 values, per phosphosite, obtained from fitting the intensity values of the 10 drug concentration points to a four-parameter logistic function.

#### I.3. Harmonized multi-omics data from the Clinical Proteomic Tumor Analysis Consortium

This dataset contains data from the *Clinical Proteomic Tumor Analysis Consortium,* which consists of multi-omics data from various cancer types. In NetworkCommons we included the proteomics, transcriptomics and phosphoproteomics data processed by the University of Michigan team’s pipeline, and then post-processed by the Baylor College of Medicine’s pipeline. Details are available in the *STAR Methods of ‘Proteogenomic Data and Resources for Pan-Cancer Analysis’* (i.e., ‘BCM pipeline for pan-cancer multi-omics data harmonization’) (Li et al. 2023).

#### I.4. Basal multi-omics data from the *NCI-60* cancer cell line panel

This dataset contains processed data from the *NCI-60* cell line panel (Shoemaker 2006), downloaded from the *CosmosR* package (Dugourd et al. 2024). It includes three files: transcriptomics, metabolomics, and the TF activities derived from the former.

#### I.5. Prior knowledge networks from OmniPath

*OmniPath* is a comprehensive collection of signaling pathways and regulatory interactions (Türei et al. 2021). It prioritizes literature curated interactions with causality information, and also provides large collections of additional kinase-substrate, ligand-receptor, signaling pathway, miRNA and gene regulatory interactions. Using the *OmniPath* API, NetworkCommons is able to import custom networks that are readily usable by its methods.

### II. Network contextualization methods

#### II.1. Reachability filter

The *reachability filter* generates a network consisting of all reachable nodes from a set of starting nodes (Fig S1, A). From a source node (blue), the *reachability filter* removes nodes that are disconnected from it, taking into account the directionality of the network. In other words, all nodes that are upstream from the source node are removed. Unlike the *all paths* algorithm, there is not a maximum depth of search.

#### II.2. Shortest (sign-consistent) paths

The *shortest path* is an algorithm for finding one or multiple paths that minimize the distance from a set of starting nodes to a set of destination nodes in a weighted graph (Dijkstra 1959) (Fig S1, B). When there are multiple paths that have the same length, then all these paths are retrieved (not to be mistaken with the *all paths* algorithm, which retrieves all paths irrespective of their length). Both *shortest paths* and *all paths* can be subject to *sign consistency* check (whether the sign of the path multiplied by the sign of the source equals the sign of the target) (Fig S1, C)

#### II.3. All (sign-consistent) paths

The *all paths* method finds all possible connections between a set of source nodes and a set of target nodes (Fig S1, D). In contrast to the *shortest path* method or the *sign consistency* method, it does not take the distance or any sign information into account, respectively. However, since this problem can exponentially explode the further away the nodes are, the function has a parameter that specifies the furthest away the algorithm is allowed to search for a path.

#### II.4. Random walk with restart (RWR) methods: Personalized PageRank

The *Personalized PageRank* (Page et al. 1999) algorithm initially calculates a weight for each node in a graph based on a *random walk with restart* method. It starts at a set of source or target nodes and determines the importance of the other nodes in the graph based on the structure of the incoming or outgoing edges. It then builds a network considering the highest-ranking nodes starting from each of the source and the target nodes. This implementation of *Personalized PageRank* follows the same philosophy as a *heat diffusion kernel* (Fig S1, E). Nodes closer to the source or target layers will receive a higher score than those further away, following the assumption that the nodes closer to the measured nodes will be more important in providing biological context compared to those far away. The nodes can be then selected using a quantile threshold (e.g ∼ N percent of the nodes with highest PPR score)

**Supplementary Figure 1:**
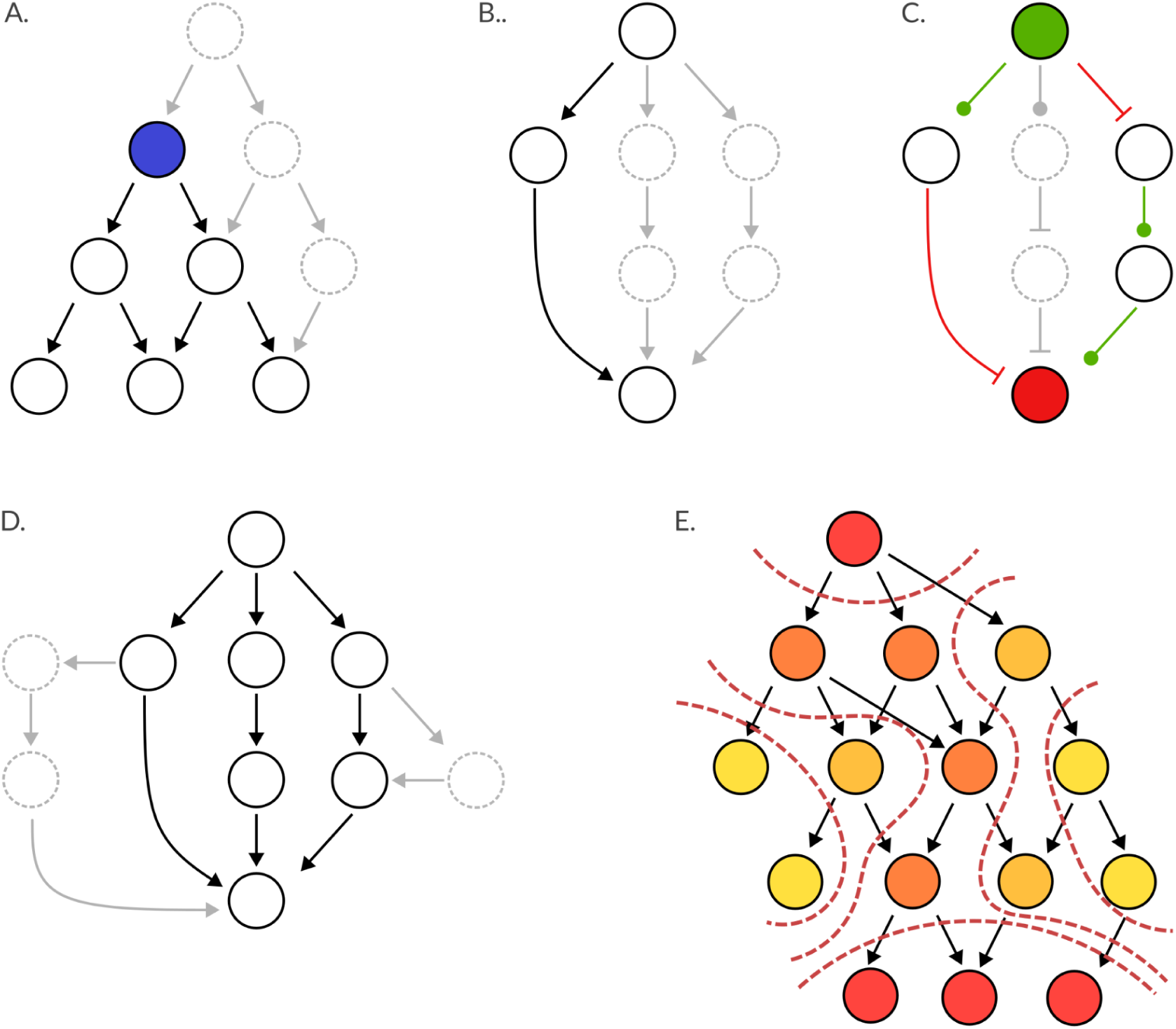
Graphical representation of topological and diffusion-like methods included in NetworkCommons: A: Reachability filtering, B: Shortest paths, C: Sign consistency check, D: All paths, E: Personalized PageRank.

#### II.5. Integer linear programming (ILP) based methods

*CORNETO (Constrained Optimization for the Reconstruction of NETworks from Omics;* (Rodriguez-Mier et al. 2024) is a unified network inference framework which combines a wide range of network methods including *CARNIVAL* (Liu et al. 2019)*CAusal Reasoning for Network identification using Integer VALue programming;* (Liu et al. 2019), which we made available in NetworkCommons. *CARNIVAL* connects a set of weighted target and source nodes using *integer linear programming* (ILP) and predicts the sign for the intermediate nodes (Fig S2, A). Thereby, it optimizes a cost function that penalizes the inclusion of edges as well as the removal of target and source nodes (Fig S2, B).

**Supplementary Figure 2:**
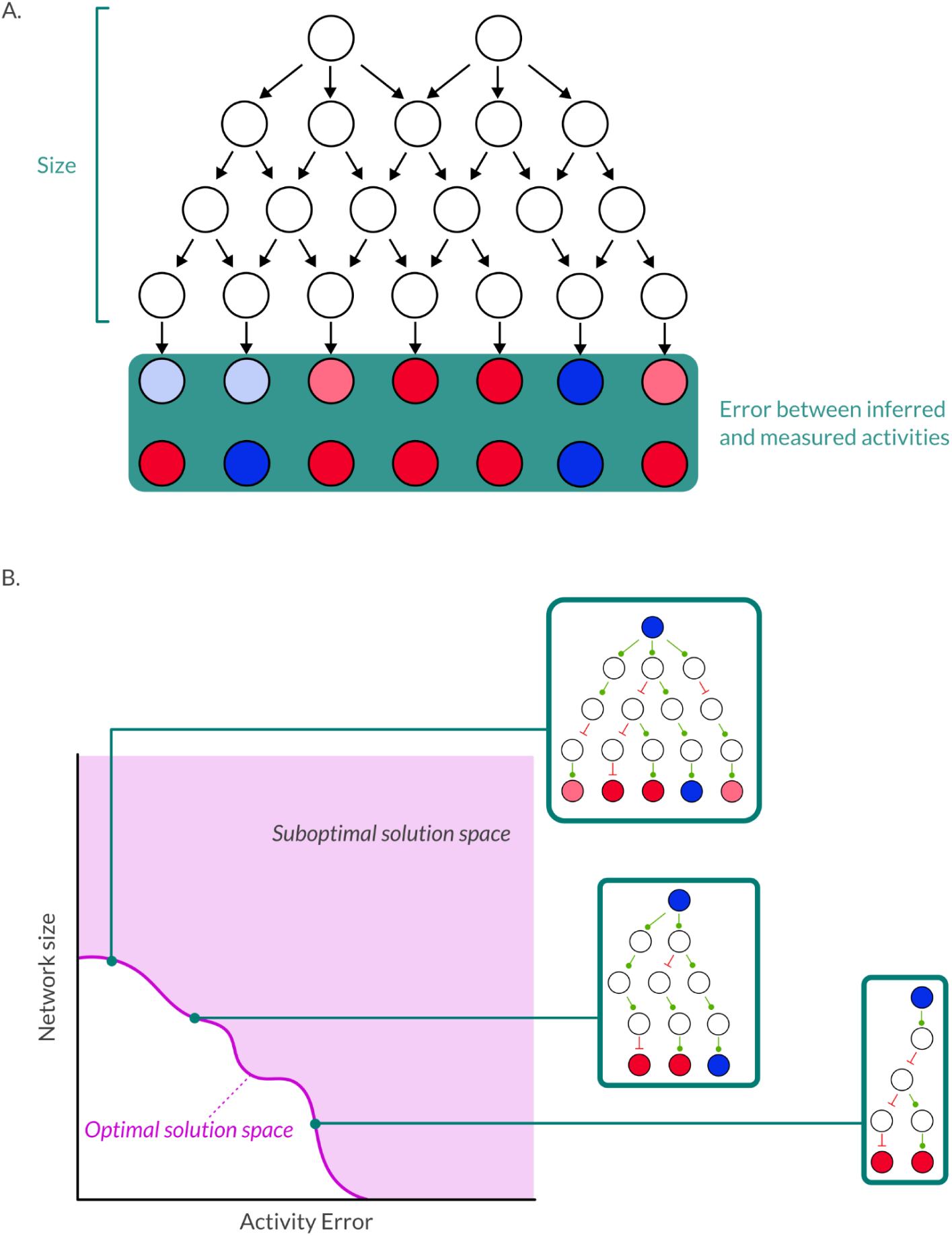
description of CARNIVAL as implemented in CORNETO. A: On the one hand, the model tries to get a sign consistent subnetwork as small as possible. In other words, the inclusion of nodes/edges is penalized. On the other hand, the model tries to minimize the difference between inferred and measured activities in downstream measurements. The regularization parameter β controls the relationship between these two factors. B: Given a PKN and a set of measurements, the interplay between size and activity error generates a family of optimal solutions (pink line) within the solution space (pink area). Changing the regularization parameter allows us to traverse this optimal solution space to find different solution networks.

#### II.6. Recursive propagation methods

The *MOON* (meta-footprint) method performs iterative footprint activity scoring and network diffusion from a set of target nodes to generate a sign consistent network (Dugourd et al. 2024); Fig S3). Starting from a set of weighted target nodes it calculates a weight for the next layer of upstream nodes using a univariate linear model. This process is repeated until a set of source nodes or a certain number of steps is reached. Hereby, any source node with an incoherent sign between *MOON* and the input sign is pruned out along with all incoming and outgoing edges. Additionally, edges between two inconsistent nodes are removed.

**Supplementary Figure 3:**
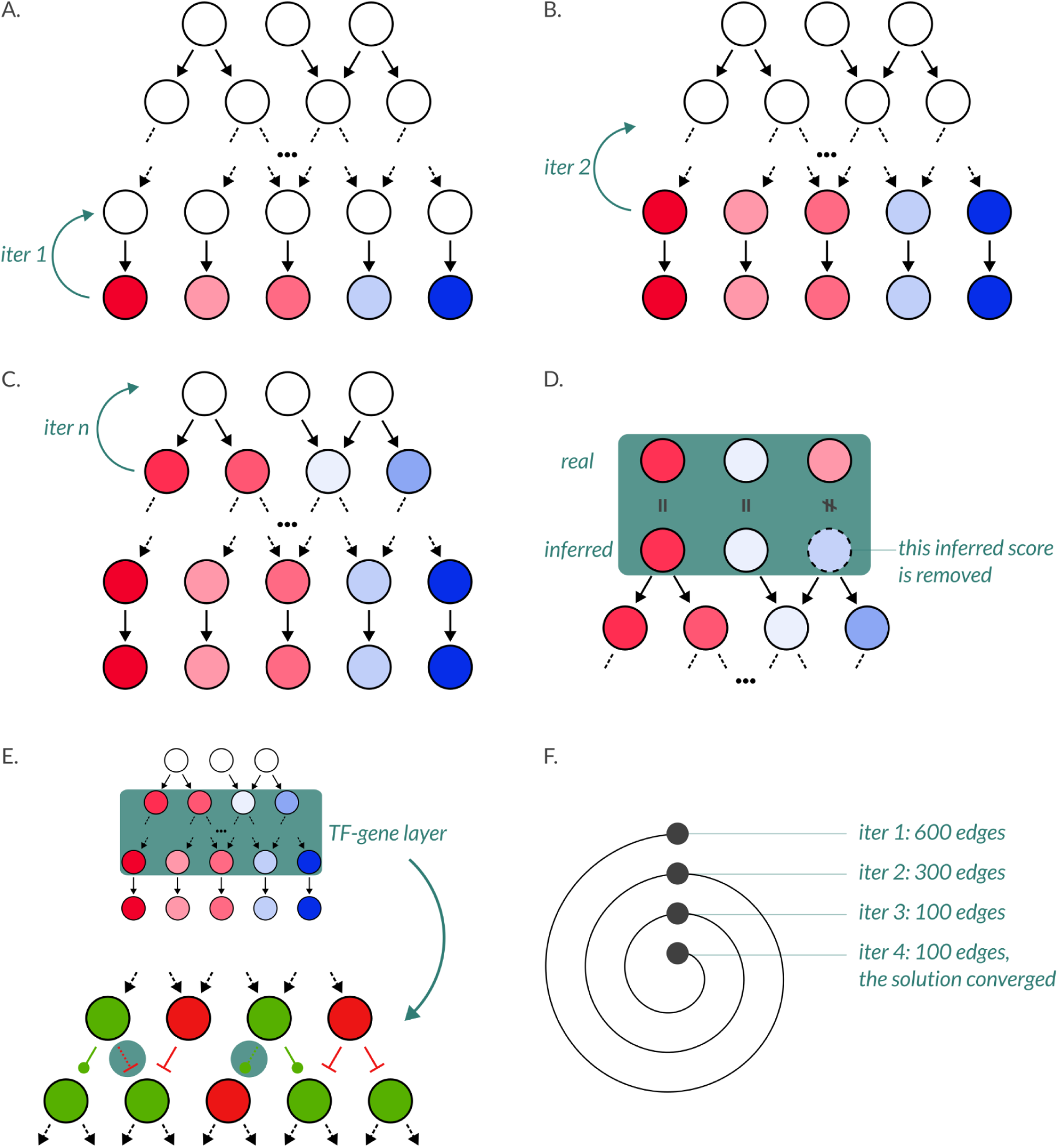
Graphical representation containing the main steps from MOON. A: The process starts with a set of downstream measurements. An univariate linear model (ulm) is used to infer the activity scores of the layer immediately upstream of the measurement layer. B: This process is then iteratively repeated in an upstream direction, computing scores for each of the nodes which have at least one child node with an associated score. C: The loop continues until the method reaches the upstream layer or a maximum number of layers have been explored. D: The MOON scores of the upstream layer are then compared against the real upstream values (if available) to check for sign consistency of the upstream layer. Otherwise, the inferred measurements for the upstream layer are removed. E: Then, the method checks the sign consistency of the TF-to-gene layer: if the sign of the TF score is not concordant with the sign of the measured gene expression + the nature of the interaction (activation or inhibition of expression), the edge between TF and gene is removed (blue circles). F: If there is any edge removal, then the process is restarted in step 1. The cycle continues until no edge is removed in step 5 (the solution has converged), or a maximum number of iterations is reached (the solution has not converged).

#### II.7. SignalingProfiler

SignalingProfiler pipeline connects source and target proteins in two steps, combining topology-based methods and CORNETO-CARNIVAL ((Venafra, Sacco, and Perfetto 2024); Fig S4). First, SignalingProfiler generates the Naïve Network, a hierarchical and multi-layered network, where each layer is defined by a set of molecular functions (Step 1). The molecular function for each protein is obtained by parsing the UNIPROT database GO Molecular Function annotation according to relative GO Ancestor terms. Then, CORNETO - CARNIVAL retrieves only sign-consistent edges from the Naïve Network (Step 2).

**Supplementary Figure 4.**
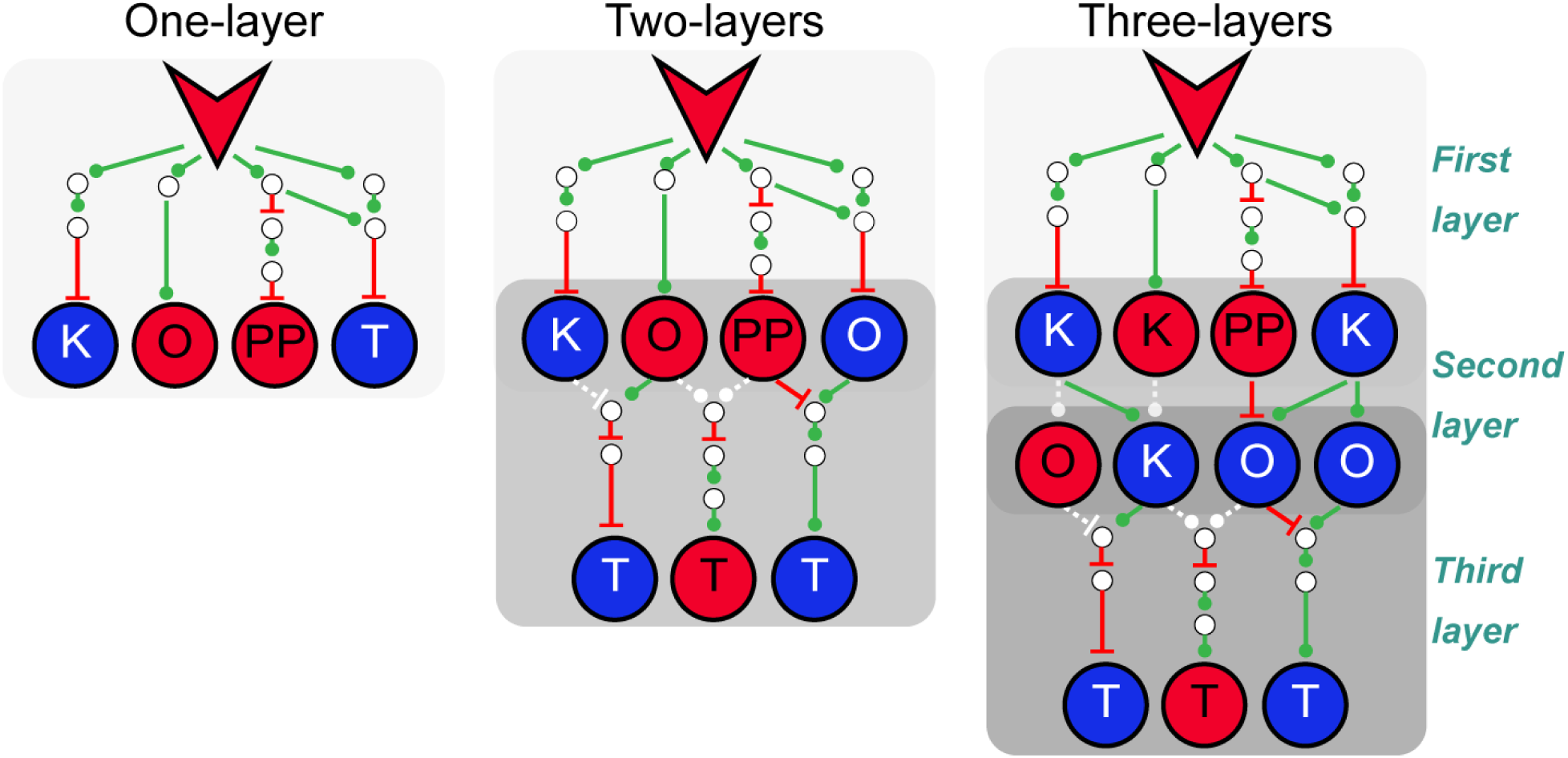
Description of SignalingProfiler. First, SignalingProfiler generates the Naïve Network, a hierarchical and multi-layered structure between source and target nodes using a topology based-method retrieving all paths of a user-specified length. Three layouts can be chosen, defined as one-, two-, or three-layers networks, with an increasing level of deepness. Each layer is defined by a different set of molecular functions referring to a signal transduction context: K, kinase; PP, phosphatases; T, transcription factor; O, all the other molecular functions. In the one-layer network, the perturbed node is connected to every target and is molecular function agnostic. The two-layers network connects the perturbed node to kinases/phosphatases/others (first layer) and then connects the latter to transcription factors (second layer). The three-layers network adds another layer between kinases/phosphatases and other signaling proteins. In the second step, SignalingProfiler calls “CORNETO - CARNIVAL” to retrieve only sign-consistent edges from the Naïve Network (removing white dashed edges).

#### II.8. Methodological categories

Because of the diverse terminology employed in the field of network biology, we link each of the methods implemented here with the methodological categories established in (Garrido-Rodriguez et al. 2022).

**Table 1.** Category of the methods according to the methodological categories described in (Garrido-Rodriguez et al. 2022).

| Method /<br>Category in<br>Garrido-Rodriguez<br>et al. 2022 | Edge filtering and<br>shortest path | Recursive<br>propagation and<br>diffusion | Integer linear<br>programming |
| --- | --- | --- | --- |
| II.1 | X |  |  |
| II.2 | X |  |  |
| II.3 | X |  |  |
| II.4 |  | X |  |
| II.5 |  |  | X |
| II.6 | X | X |  |
| II.7 | X |  | X |

### III. Evaluation strategies

#### III.1. Pathway enrichment analysis

We use the nodes of the context specific subnetworks to perform *overrepresentation analysis* (ORA) against a set of predefined gene sets, among which we expect one to be especially represented (for example, a specific pathway will be overrepresented if said pathway is perturbed, or is especially active/inactive in a given profile). Therefore, the methods producing networks in which the selected geneset(s) is ranked high, according to their ORA score, against others, will have a better performance (Fig S5).

**Supplementary Figure 5:**
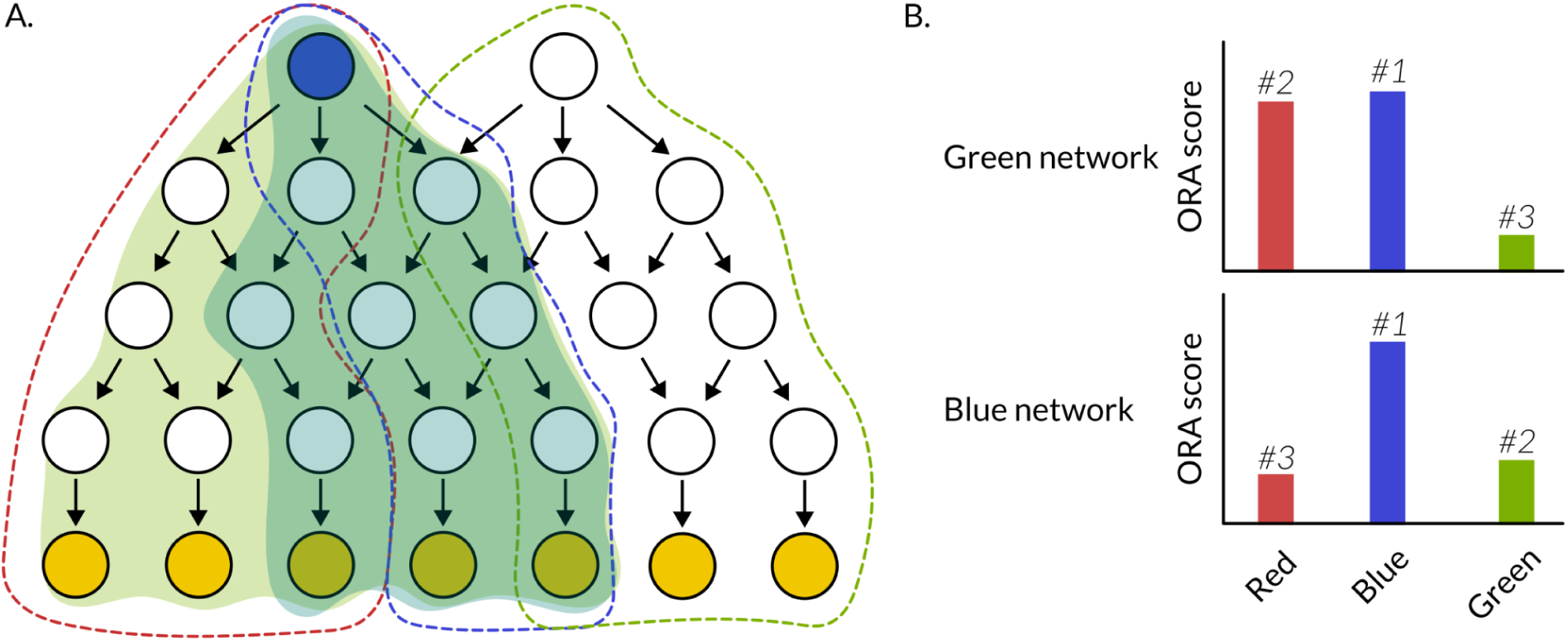
Evaluation strategy based on pathway enrichment. Given a set of predefined pathways (A), a good performance is reached when the ORA score of the perturbed pathways is higher than the ORA score of those not perturbed (B).

#### III.2. Drug perturbation off-target recovery

In this setting, we assume that, in the context of a drug perturbation, the effects measured via omics data are not only a product of the perturbation origin (the drug’s main targets), but also of secondary downstream effectors or drug off-targets. Therefore, in order to contextualize the perturbation, we expect that these drug off-targets are included in the subnetwork, too. We used the gold standard from the PANACEA DREAM challenge (Douglass et al. 2022), which provides drug-target links. Methods that recover a higher share of off-targets, compared to a random control, will be more successful in contextualizing the perturbation, since the method incorporates the off-targets’ effect (Fig S6).

**Supplementary Figure 6:**
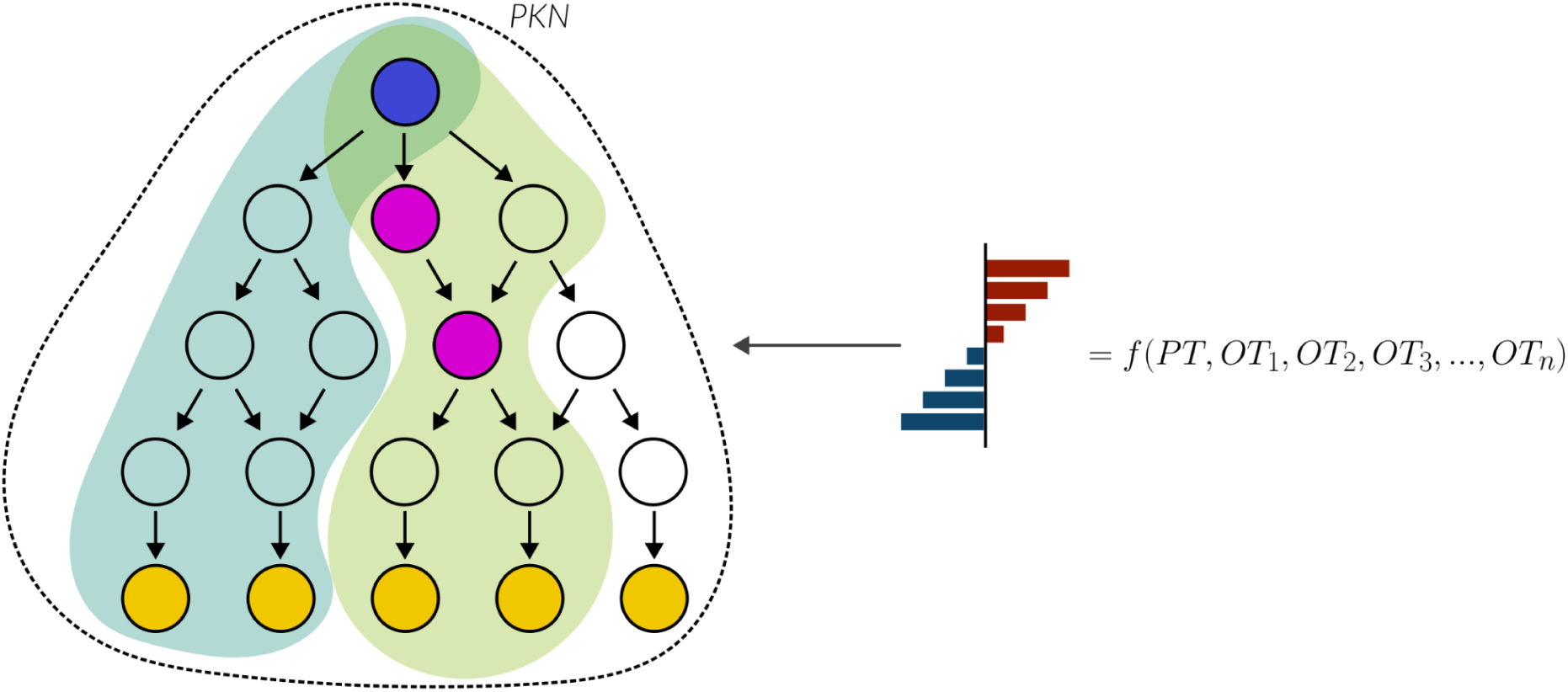
Depiction of the off-target recovery strategy. Good performance is reached when all off-targets of a drug are recovered in the solution network.

#### III.3. Sensitive phosphoproteins recovery

In this setting, we assume that, in a perturbational context, those elements in a network that are more sensitive to a perturbation (have a lower EC50) will be more important in the contextualisation of said perturbation. Now, we compare these results to a set of random controls. To build these random controls, we shuffle the measurement labels while leaving the network topology intact. In this case, we would expect that, since the measurements no longer codify any perturbation, the resulting networks would make no sense biologically. Therefore, the differences in sensitivity between nodes included and excluded from the network would be minimal. Methods producing result networks whose nodes have a low average EC50 (compared to nodes not included in the network) are better performers than those producing networks where this difference (EC50_in - EC50_out) is not that big (Fig S7).

**Supplementary Figure 7:**
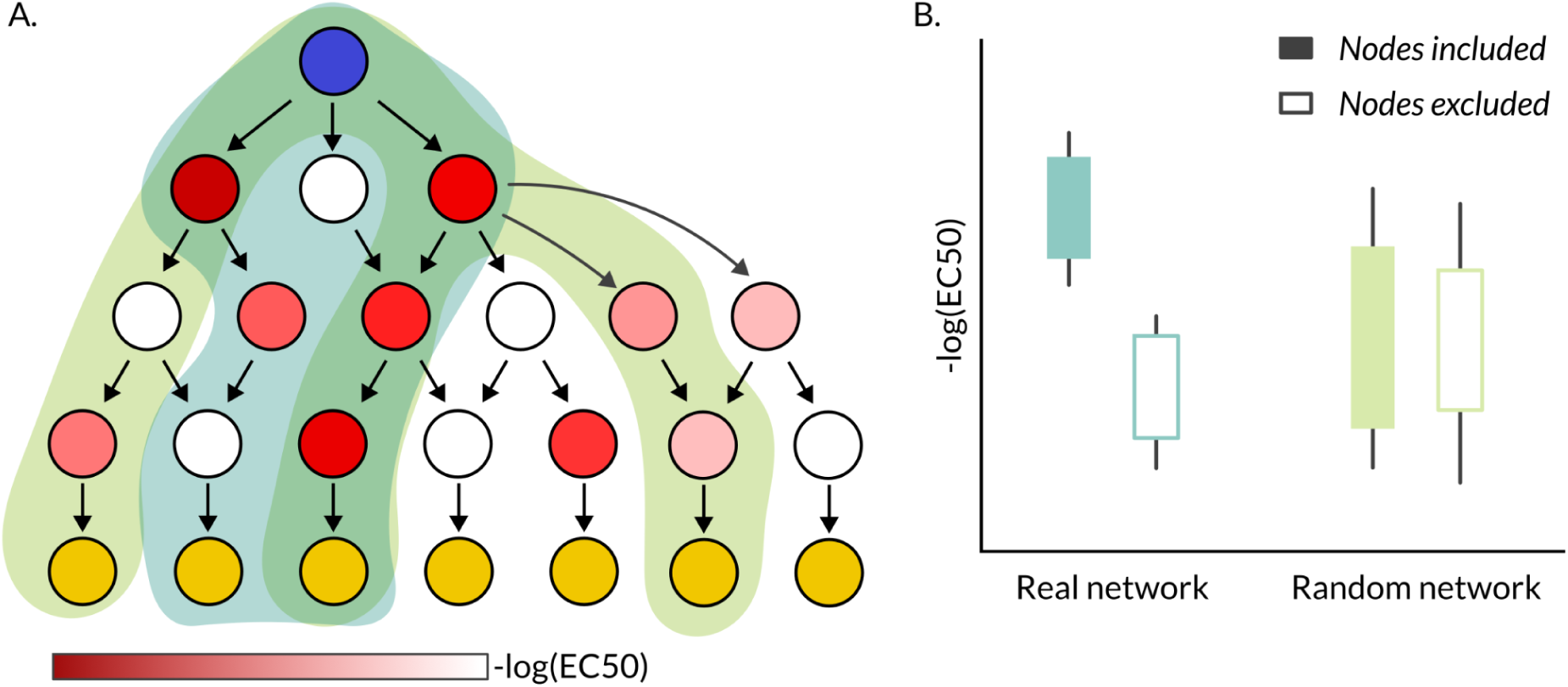
A: Two networks, a real one (blue) and a random one (green) are contextualized from a PKN. B: Using sensitivity data (EC50 values), a good-performing method should return solution networks which are enriched in sensitive elements (the -log(EC50) should be higher in the nodes inside the solution network compared to the nodes excluded from it. To establish a baseline, random controls are also performed, in which the expected result would be no differential enrichment in sensitive nodes in the solution network compared to the overall PKN.

#### III.4. Multi-omics validation: Recovery of dysregulated kinases

This approach makes use of three different types of omics: proteomics, phosphoproteomics and transcriptomics. Using proteomics, we determine the most abundant (as a proxy for active) receptors. Using transcriptomics, we compute TF enrichment analysis and select the ones with the highest absolute activity score. Then, using the receptors as upstream nodes and the TFs as downstream nodes, we can obtain contextualized prior-knowledge networks. For the evaluation, we can use the phosphoproteomics data to compute kinase activities. These activities can be used to evaluate whether the solution networks contain a higher amount of dysregulated kinases. In other words, the methods should maximize the difference between average kinase activity of the nodes returned in the solution network, compared to the average in the overall prior-knowledge network. Besides this, the difference should also be higher than the one between a random control and the PKN (Fig S8).

**Supplementary Figure 8:**
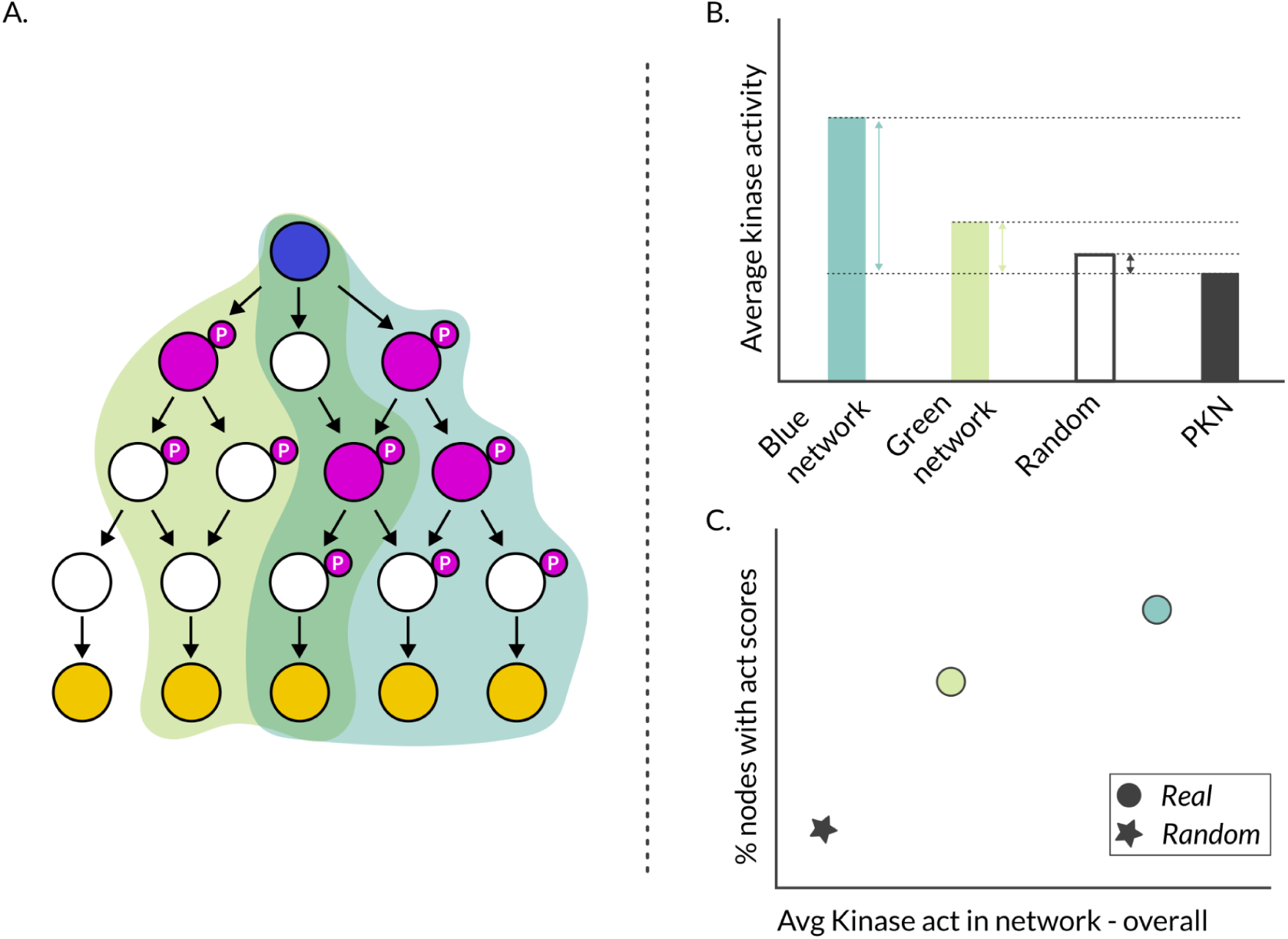
A: A network, containing kinases (purple), is contextualized using different methods which return different solution networks (blue, green). B: by inferring kinase activities for the recovered kinases, using the phosphorylation status of their targets, we obtain an average kinase activity value per network. C: Comparing this value versus the overall activity taking into account all kinases in the PKN, we can determine how enriched the solution networks are in active kinases, compared to a random control. A good performer would achieve both high coverage (% of recovered nodes being kinases, gives information about robustness of the score) and high difference in activity compared to the PKN (gives information about performance).

### IV. Implementation, API and contribution

Table 2. lists the main functionalities incorporated from dependencies used in NetworkCommons with the third party packages used for the implementation.

**Table 2:** Overview of the most important algorithms from other packages used in NetworkCommons.

| Module | Function-NC | Function-pkg | Backend | Version |
| --- | --- | --- | --- | --- |
| data.network | get_omnipath | interactions.AllInteractions.get | OmniPath | 1.0.8 |
| data.omics | get_ensembl_mappings | biomart.BiomartServer | biomart | 0.9.2 |
|  | deseq2 | _deseq2_default_inference.DefaultInference | pydeseq2 | 0.4.11 |
|  |  | default_inference.DeseqDataSet | pydeseq2 | 0.4.11 |
|  |  | dds.deseq2 | pydeseq2 | 0.4.11 |
|  |  | ds.DeseqStats | pydeseq2 | 0.4.11 |
| eval | run_ora | get_ora_df | decoupler-py | 1.8.0 |
| methods | run_corneto_carnival | methods.runVanillaCarnival | corneto | 1.0.0.dev0 |
|  |  | methods.carnival.get_selected_edges | corneto | 1.0.0.dev0 |
|  | run_shortest_paths | all_shortest_paths | networkx | 3.2.1 |
|  | run_reachability_filter | descendants | networkx | 3.2.1 |
|  | compute_all_paths | all_simple_paths | networkx | 3.2.1 |
|  | add_pagerank_scores | pagerank | networkx | 3.2.1 |
|  | run_moon_core | run_wmean | decoupler-py | 1.8.0 |
|  | run_moon_core | run_ulm | decoupler-py | 1.8.0 |
|  | get_ego_graph | ego_graph | networkx | 3.2.1 |

#### IV.1. Prior knowledge

Most of the methods in NetworkCommons operate on *NetworkX Graph* objects, and the easiest way to include custom prior knowledge or new resources is to convert them into a *NetworkX* object. Such conversion code can be included in the `data.network` module of NetworkCommons, making the resource available for all users of the platform. The *Graph* object from *NetworkX* is interconvertible with the *Graph* object provided by *CORNETO,* the required input for some of the methods. In the future we plan to wrap these third party graph objects into our own *Network* object, providing more features and tighter integration.

#### IV.2. Omics data

The `data.omics` module provides a simple caching downloader and a minimal API to download and load datasets from repositories. We established a directory at https://commons.omnipathdb.org/ where we host some of the currently built-in datasets, and are happy to include datasets from contributors. Any other service accessible by HTTP is suitable. Loaders for datasets can be implemented in an analogous way to the existing ones in the `data.omics` module. Each loader returns one or more `pandas` data frames. We also provide integration to *DeSeq2* (Muzellec et al. 2023) for differential expression analysis. See more details in the *Contribution guidelines* section of the documentation.

#### IV.3. Methods

The network contextualization methods, described in section II. of this Supplementary Data, are implemented in the `methods` module of NetworkCommons. We created a functional API, where each method consists of one or more functions within its own submodule, which accept Network objects, omics data and parameters, and implement the inference workflow. New methods can be added in a similar way to existing ones, as we describe in detail in the *Contribution guidelines* section of the documentation. For methods already implemented in other packages, we recommend creating a minimal wrapper in the `methods` module, for seamless integration. In the future, based on initial feedback and experiences, we plan to provide higher level objects for further integration of the APIs and workflows.

#### IV.4. Evaluation

The methods for evaluation of context specific networks, described in section III. above, are located in the `eval` module of NetworkCommons. The functional API in this module provides a set of indicators of performance, according to the evaluation strategies detailed in the documentation. Many of these functions are reusable when implementing new evaluation strategies. See more details in the *Contribution guidelines* section of the documentation about implementing new strategies, and about the current ones in the *Details* section. For practical examples, please check the *Vignettes* section.

## 5. Acknowledgements

We would like to thank Aurelien Dugourd, Benjamin Maier, Evangelia Petsalaki and Laurence Calzone for the inspiring discussions that contributed to our design. DT and OI were supported by the HPC/Exascale Centre of Excellence for Personalised Medicine in Europe [PerMedCoE; European Union Horizon 2020 program, grant no. 951773], DT also by the German Federal Ministry of Education and Research [Bundesministerium für Bildung und Forschung BMBF; grant no. 031L0181B] funds granted to JSR, and OI by the German Research Foundation (DFG) grant 411368829 SA 3536/2-1. SMD was funded by the German Federal Ministry of Education and Research, particularly the LiSyM Cancer research core (BMBF, Funding number: 031L0257B). PRM was supported by DECIDER [European Union Horizon 2020 program, grant no. 965193]. MGR was supported through state funds approved by the State Parliament of Baden-Württemberg for the Innovation Campus Health + Life Science Alliance Heidelberg Mannheim.

## 7. Authors contributions

VPG, DT, LP, MGR and JSR: Conceptualization; VPG, DT, OI, VV: Software; VPG, PRM, SMD, MGR and JSR: Methodology; VPG, OI: Visualization. VPG, DT, MGR and JSR Writing – Original Draft Preparation. MGR, JSR: Project Administration. JSR: Funding Acquisition.

